# Endosymbiotic adaptations in three new bacterial species associated with *Dictyostelium discoideum*: *Burkholderia agricolaris* sp. nov., *Burkholderia hayleyella* sp. nov., and *Burkholderia bonniea* sp. nov

**DOI:** 10.1101/304352

**Authors:** Debra A. Brock, Alicia N.M. Hubert, Suegene Noh, Susanne DiSalvo, Katherine S. Geist, Tamara Haselkorn, David C. Queller, Joan E. Strassmann

## Abstract

Here we name three species of *Burkholderia* that can defeat the mechanisms by which bacteria are normally excluded from the spores of a soil dwelling eukaryote *Dictyostelium discoideum*, which is predatory on bacteria. They are *B. agricolaris* sp. nov., *B. hayleyella* sp. nov., and *B. bonniea* sp. nov. These new species are widespread across the eastern USA and were isolated as internal symbionts of wild collected *D. discoideum*. Evidence that they are each a distinct new species comes from their phylogenetic position, carbon usage, reduced cell length, cooler optimal growth temperature, and ability to invade *D. discoideum* amoebae and remain there for generations.

## Importance

The evolutionary origins of symbioses are best investigated in systems that retain flexibility in association. In these interactions, the tensions between host and symbiont will be more dynamic and conflicts more easily assessed. One recently developed example is the symbiosis between the social amoeba *Dictyostelium discoideum* and the three new species of *Burkholderia* presented here. All three of these new species facilitate the prolonged carriage of food bacteria by *Dictyostelium discoideum*. Further studies of the interactions of these three new species with *D. discoideum* should be very fruitful for understanding the ecology and evolution of mutualistic symbioses.

## Introduction

We are only beginning to appreciate that every feature of eukaryotes has evolved in a microbial world (1). Eukaryote soil-dwelling amoebae are particularly exposed to bacteria in their environment. They may be penetrated by bacteria using secretion systems (2). They may ingest bacteria that foil their digestive systems and take up residence inside their cells (3). Some bacteria may become permanent or semi-permanent residents (4). In this study, we examine the characteristics of *Burkholderia* that have formed symbiotic relationships with the social amoeba *Dictyostelium discoideum* (4). Based on data presented here and data previously published, we name three new species in the plant beneficial clade of *Burkholderia*.

The genus *Burkholderia* is comprised of over 60 species that were originally included in the genus *Pseudomonas*, but was identified as unique by Yabuuchi et al. in 1992 (5). *Burkholderia* are diverse and include species that are adapted for life in the soil, as endosymbionts, and as pathogens for both plants and animals. Of the pathogenic bacteria, there is a group of 18 species that are together identified as the *Burkholderia cepacia* complex (BCC), which are most predominantly associated with infections that can be lethal in immunocompromised human patients, most notably, patients with cystic fibrosis (6). Because of the wide variety of species Sawana et al. (7) proposed separating the genus into two separate genera: *Burkholderia*, which contains the pathogenic species and *Paraburkholderia* which contains the environmental species. However, further examination of this clade reveals that this separation and reclassification is premature because of difficulties in placing intermediate species (8). Therefore, we stick with the original genus name, *Burkholderia*.

To support naming new species, we examined multiple isolates of each species in several ways. First, we have already established that they can cause the farming trait in *D. discoideum*, where farming is the ability to carry food bacteria through the social stage and then release and consume it after the spores hatch (4). Second, we place the *Burkholderia* isolates in a phylogeny along with other *Burkholderia* species. Third, we examine carbon usage using a suite of possible carbon food sources. Fourth, we measure the length of the bacterial cells. Fifth, we investigate optimal growing temperatures. Based on these data, we name 3 new species.

## Results

*B. agricolaris* sp. nov., *B. hayleyella* sp. nov., and *B. bonniea* sp. nov. are gram-negative, motile, rod-shaped, β-proteobacteria. We isolated these bacteria in association with field-collected clones of *D. discoideum* (Table 1). These symbiotic species can be differentiated from each other and from other *Burkholderia* by their endosymbiotic habit (4), phylogenetic placement, carbon usage profile, cell length, and optimal temperature range.

**Table 1.** Summary table of comparisons for all species, including a subset of carbon usage types. Included in this table are the number of *Burkholderia* sp. nov. isolates in each group, whether or not the isolate is associated with *D. discoideum*, average bacterial length, and optimal growth temperature. Next, a subset of about one-third of the relevant carbon types follows. A plus symbol indicates all isolates were able to use a specific carbon and a minus symbol indicates the opposite. A number value indicates the percentage of isolates in a specific group that can utilize that particular carbon. *B. agricolaris* and *B bonniea* are able to utilize some carbons that their close *Burkholderia* relatives cannot.

| Characteristic | *B.*<br>*fungorum*<br>ATCC<br>BAA-463 | *B.*<br>*xenovorans*<br>LB400 | *B.*<br>*phymatum*<br>STM-815 | *B.*<br>*agricolaris*<br>sp. nov | *B.*<br>*hayleyella*<br>sp. nov | *B.*<br>*bonniea*<br>sp. nov |
| --- | --- | --- | --- | --- | --- | --- |
| n (# strains) | 1 | 1 | 1 | 7 | 7 | 2 |
| *Dictyostelium* | - | - | - | + | + | + |
| Cell length (µm) | 1.57 ± 0.03 | 1.66 ± 0.03 | 1.7 ± 0.03 | 1.45 ± 0.009 | 1.36 ± 0.009 | 1.38 ± 0.014 |
| Optimal growth temperature (°C) | 37 | 30 and 37 | 37 | 30 | 30 | 30 |
| Maltose | - | - | - | 43% | - | - |
| D-Cellobiose | - | - | - | 71% | - | - |
| α-D-Lactose | - | - | - | 71% | - | - |
| γ-Hydroxy Butyric | - | + | - | 57% | - | - |
| Inosine | - | - | - | 86% | - | + |
| D-Melibiose | + | - | - | - | - | - |
| Xylitol | + | - | + | 57% | - | - |
| Glycyl-L-Aspartic | + | + | - | 71% | - | - |
| Gentiobiose | + | + | - | 57% | - | 50% |
| Glucuronamide | + | + | + | 57% | - | - |
| α-Keto Valeric Acid | + | + | + | 71% | - | - |
| 2-Aminoethanol | + | + | + | 71% | - | - |
| D-Galacturonic Acid | + | + | + | 86% | - | - |
| L-Fucose | + | + | + | + | 29% | - |
| D-Galactose | + | + | + | + | - | 50% |
| Mono-Methyl- | + | + | + | + | - | 50% |
| N-Acetyl-D- | + | + | + | + | - | - |
| L-Arabinose | + | + | + | + | - | - |
| D-Arabitol | + | + | + | + | - | - |
| m-Inositol | + | + | + | + | - | - |
| D-Mannitol | + | + | + | + | - | - |
| L-Rhamnose | + | + | + | + | - | - |
| D-Sorbitol | + | + | + | + | - | - |
| Formic Acid | + | + | + | + | - | - |
| D-Galactonic Acid | + | + | + | + | - | - |
| D-Glucuronic Acid | + | + | + | + | - | - |
| Malonic Acid | + | + | + | + | - | - |
| Quinic Acid | + | + | + | + | - | - |
| D-Saccharic Acid | + | + | + | + | - | - |
| Sebacic Acid | + | + | + | + | - | - |
| L-Phenylalanine | + | + | + | + | - | - |
| L-Pyroglutamic Acid | + | + | + | + | - | - |

### Phylogenetic analysis

The phylogeny that we constructed from whole genome k-mers of 23 bases is very robust, with every node obtaining 100% bootstrap support (Figure 1). There is some structure within *B. agricolaris*. Its closest relative is *B. fungorum* strain BAA-463 from the American Tissue Culture Collection. The other two new species, *B. hayleyella*, and *B. bonniea*, are each other’s closest relatives and their long-branch lengths indicate they are quite diverged from anything else in this phylogeny. The sister taxon to *B. hayleyella* and *B. bonniea* is the entire clade containing *B. agricolaris*.

**Figure 1.**
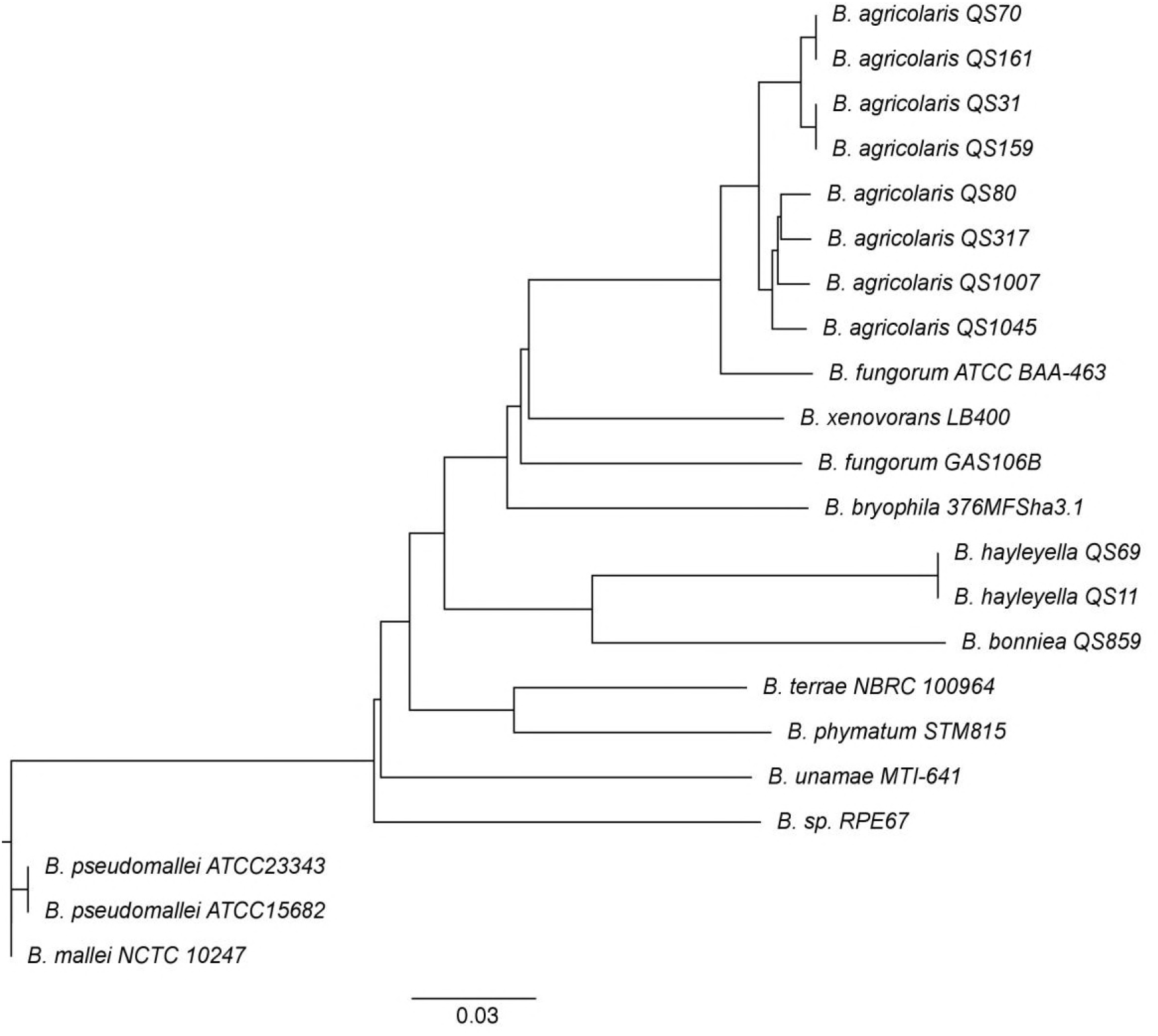
Phylogeny of three new non-pathogenic *Burkholderia* species found in association with *D. discoideum*. We estimated the phylogenetic relationships of these symbionts with 11 known plant-associated and environmental *Burkholderia* species. We used k-mers frequencies (k=23) of each genome and created an alignment and assembly-free distance tree. We ran 999 nonparametric bootstraps by resampling 1/k of the rows of the shared k-mer table and found that each branching event was fully supported (bootstrap value = 1). The final tree is shown here rooted by the *B. mallei* and *B. pseudomallei* clade part of the pathogenic *Burkholderia* cluster.

### Carbon usage for *B. hayleyella* and *B. bonniea* is greatly reduced

The 95 carbon types from Biolog GN2 test plates are organized into eleven functional groups (Supplementary Table 2), and we subjected these groups to principal component analysis (PCA). The generalized output of the PCA revealed that most of the variation could be described by PC1 (79.3%) and PC2 (8.6%). PC1 is composed of ten of the eleven components and positively correlates roughly equally with nine of these components (carbohydrates, esters, carboxylic acids, amides, amino acids, aromatic chemicals, amines, alcohols, and phosphorylated chemicals) suggesting these nine criteria vary together, with a tendency towards loss in *B. hayleyella* and *B. bonniea*. PC2 is composed of six components and positively correlates strongly with two (phosphorylated chemicals and amines). One component of the PCA analysis, brominated chemicals, was utilized by all of the bacteria so was not useful in distinguishing between the carbon groups. The scatter plot of the component scores for PC1 and PC2 shows that the non-symbiotic species and *B. agricolaris* sp. nov. can use a broader range of carbon sources than *B. hayleyella* or *B. bonniea* (Figure 2). We found the most variance in carbon use was explained by an interaction between *Burkholderia* species and carbon type (χ^2^ = 497.43, DF = 43, *P* << 0.001, ΔAIC = −175.0). We also found an overall effect of both additive terms in the model, *Burkholderia* clade (χ^2^ = 58.54, DF = 3, *P* << 0.001, ΔAIC = −52.5) and carbon type (χ^2^ = 383.94, DF = 10, *P* << 0.001, ΔAIC = −363.9), over the null model. From our model, we found *B. agricolaris, B. hayleyella*, and *B. bonniea* are all significantly different from one another (Benjamini-Hochberg adjusted p-values: *B. agricolaris/B. hayleyella P* << 0.001, *B. agricolaris/B. bonniea P* << 0.001, *B. hayleyella/B. bonniea P* = 0.003). Specific differences can be found in the species descriptions.

**Figure 2.**
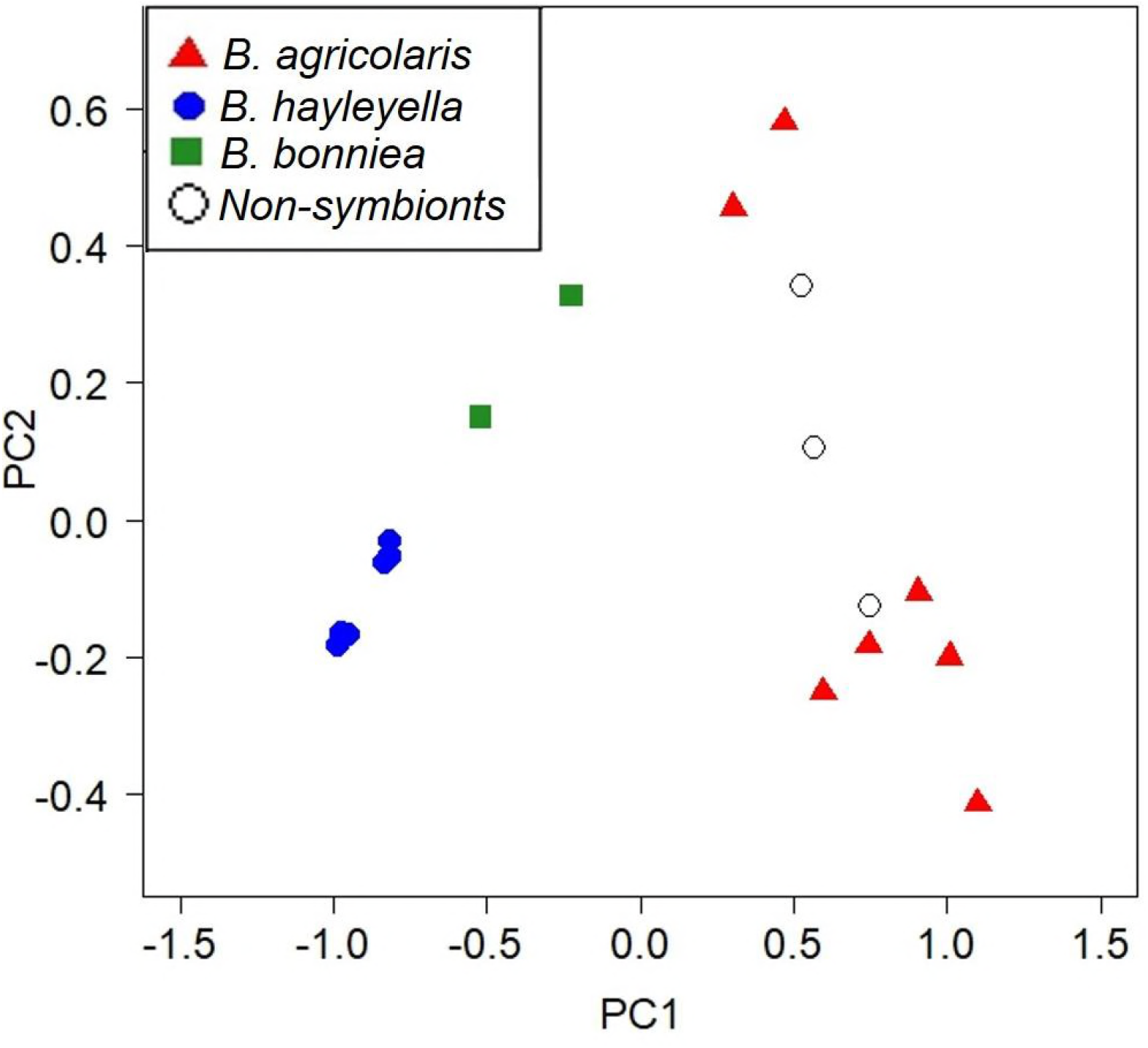
Principal component analysis plot of carbon usage. Principal Component 1 (x-axis) accounts for 79.3% of the variance and Principal Component 2 (y-axis) accounts for 8.6% of the variance. Each symbol represents one bacteria isolate with different symbols representing each species. A higher value on the x-axis represents a larger number of carbon sources that can be utilized; *B. hayleyella* and *B. bonniea* have greatly reduced carbon usage compared to *B. agricolaris* and the non-symbionts.

We also found *B. hayleyella* and *B. bonniea* are significantly different in overall carbon usage compared to the three closely related non-symbiont species while *B. agricolaris* are not (Benjamini-Hochberg adjusted p-values: *B. agricolaris* p = 0.34, *B. hayleyella P* << 0.001, *B. bonniea P* << 0.001). However, though carbon usage of *B. agricolaris* is not statistically different from the non-symbiont species, Table 1 and Supplementary Table 6 highlight differences in usage of specific carbons by *B. agricolaris* sp. nov. These include no utilization of D-melibiose by *B. agricolaris* sp.nov. compared to close sister non-symbiont *B. fungorum*, and no utilization of adonitol by *B. agricolaris* sp.nov. compared to close sister non-symbiont *B. xenovorans*. Additionally, many or all of *B. agricolaris* sp.nov. are able to utilize several carbon sources that *B. fungorum* ATCC_BAA-463 cannot. Examples are: 100% *B. agricolaris* sp.nov. tested can utilize hydroxy-L-proline, 86% tested can utilize L-ornithine and inosine, and 71% tested can utilize glycogen, D-cellobiose, and α-D-lactone.

We also found differences in the overall pattern of carbon use when we performed pairwise comparisons between species for each of the individual carbon source types (Supplementary Table 3). From these pairwise carbon use comparisons, some patterns emerge. The divergence in carbon use for *B. hayleyella* and *B. bonniea* from the non-symbiont species and *B. agricolaris* is predominantly in their use of amino acids, carbohydrates, and carboxylic acids. However, *B. hayleyella* and *B. bonniea* diverge from each other in their use of amides and carbohydrates. *B. agricolaris* and *B. hayleyella* also diverge in their use of amides. *B. agricolaris* has maintained a similar carbon use pattern to the outgroup species, but it seems to be beginning to diverge in its use of aromatic chemicals, specifically inosine.

### Symbiont bacteria length is shorter

We examined morphological differences between the new *Burkholderia* species and non-symbiont controls by measuring bacterial length (Figure 3). The generalized linear mixed model showed an overall difference in the effect of *Burkholderia* clade on length (χ^2^ = 17.19, DF = 3, *P* < 0.001, ΔAIC = −11). Between species, we found that all *Burkholderia* sp. nov. are significantly shorter in length than the outgroup species (Benjamini-Hochberg adjusted P-values: *B. agricolaris P* = 0.0013, *B. hayleyella P* << 0.001, *B. bonniea P* = 0.0012). We also found that lengths differed between *B. agricolaris* and *B. hayleyella* (*P* = 0.033), but not between *B. agricolaris* and *B. bonniea (P* = 0.32) or between *B. hayleyella* and *B. bonniea* (P = 0.67).

**Figure 3.**
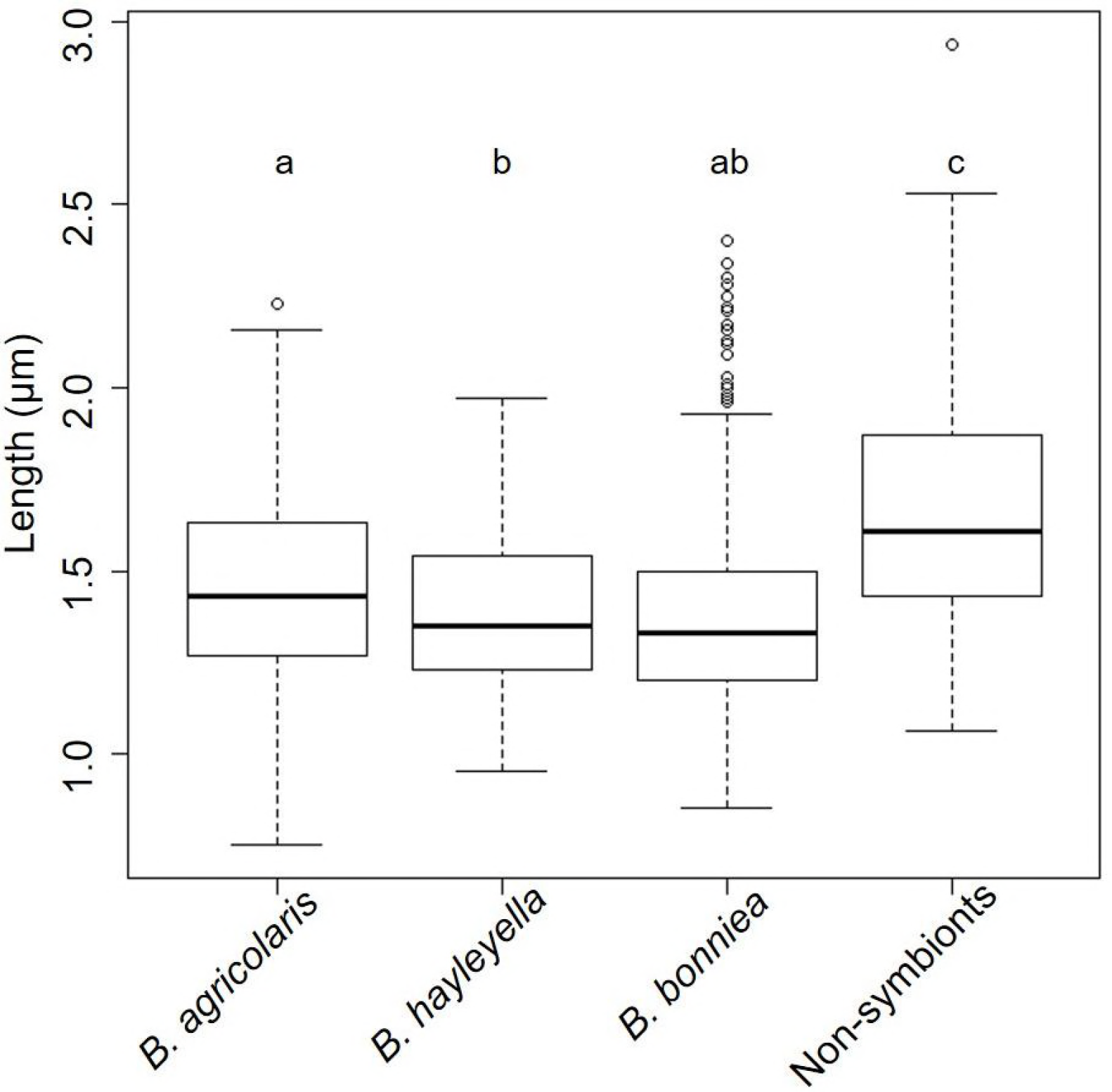
Symbiont *Burkholderia* bacteria lengths are shorter than non-symbionts. We measured the length of about one hundred bacteria for each *Burkholderia* sp. nov. and for the non-symbionts (see Supplementary table x). We used seven strains for *B. agricolaris*, seven strains for *B. hayleyella*, two strains for *B. bonniea*, and three strains for non-symbionts. We found all three symbiont bacteria species are shorter than non-symbiont bacteria species. Significant differences in length found between bacteria are indicated by different letters which reflect results of a Benjamini-Hochberg correction for multiple comparisons.

### Optimal growth temperature for symbiont *Burkholderia* sp. nov. is reduced compared to close relative non-symbiont *Burkholderia*

We tested all isolates to determine the range of temperatures permissive for growth and the optimal growth temperature. We used a range of five temperatures: 4°C, 22°C, 30°C, 37°C, and 45°C. For range of growth, we first found that none of our isolates including the non-symbiont controls were able to sustain visible growth at either 4°C or 45°C (Figure 4). The optimal growth temperature for the three symbiont *Burkholderia* sp. nov. is 30°C compared to the non-symbiont optimum of 37°C. We found all isolates of *B. hayleyella* and *B. bonniea* grew less densely overall and had no growth at 37°C compared to non-symbiont controls and *B. agricolaris. B. agricolaris* grew vigorously at 30°C and moderately well at 37°C compared to non-symbiont controls that had excellent growth at a higher optimal temperature of 37°C. We performed a Fisher’s Exact Test comparing the 22°C, 30°C, and 37°C data and excluding the 4°C and 45°C because neither symbiont nor non-symbiont isolates grew at these two temperatures. The results of the exact contingency table test (Supplementary Table 5) showed that the growth range and extent of growth differed among the four groups of *Burkholderia* (*P* < 0.001). We did post hoc tests to determine specific growth differences. Using a Bonferroni corrected cutoff of 0.008, *B. hayleyella* is significantly different from *B. agricolaris* (P = 0.000022) and the non-symbionts (P = 0.0032). *B. bonniea* is different from *B. agricolaris* (P = 0.0072), but not the non-symbionts (P = 0.063), or from *B. hayleyella* (P = 0.074). Lastly, *B. agricolaris* does not differ from the non-symbionts (P = 0.091).

**Figure 4.**
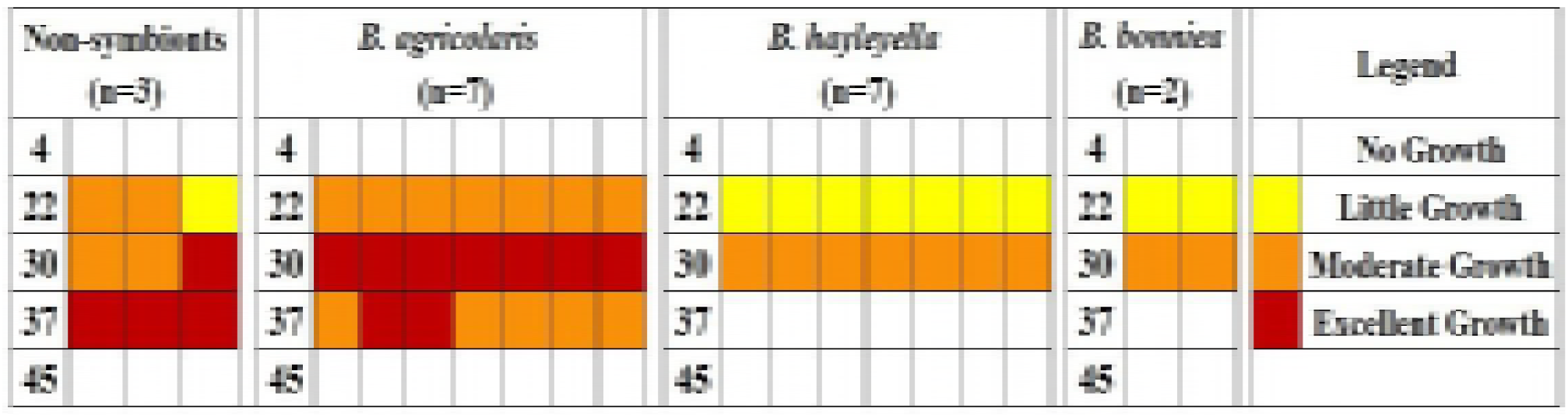
The optimal growth temperature of *Burkholderia* sp. nov. is 30°C. We tested a range of temperatures to determine growth range and optimal temperature. *B. hayleyella* and *B. bonniea* have reduced range of growth and grow less densely compared to *B. agricolaris* and the non-symbionts. *B. agricolaris* has the same range as the three non-symbiont *Burkholderia* but grows less densely at 37°C and best at 30°C.

### Description of *Burkholderia agricolaris* sp. nov., *Burkholderia hayleyella* sp. nov., and *Burkholderia bonniea* sp. nov

*Burkholderia agricolaris* (uh´gri.ko.la.ris L. fem. adj. *agricolaris* facilitating farming). The morphology of colonies is off-white, domed, and shiny with smooth edges. Bacteria are motile, non-sporulating, straight rods. The G+C content varies between 61·5 and 61·9 mol% calculated from whole genomic sequences. The strains are stored in a sterile 20% glycerol solution at −80°C and subcultured on SM/5 agar plates at 22°C. We isolated the type strain BaQS159 as a symbiont of wild *D. discoideum* clone QS159 in May 2008. *D. discoideum* QS159 was isolated from soil and leaf litter collected from Mountain Lake Biological Station in April 2008. The G+C content of the type strain is 61·6 mol% calculated from the whole genomic sequence.

Good growth at 30°C, weak growth at 22°C and 37°C, and no growth at either 4°C or 45°C on nutrient agar containing glucose, bactopeptone, and yeast extract. The type strain BaQS159 has the ability to utilize the following carbon sources as determined by the Biolog GN2 test panel: Tween 40;Tween 80; N-Acetyl-D-galactosamine; N-Acetyl-D-glucosamine; Adonitol; L-Arabinose; D-Arabitol; D-Fructose; L-Fucose; D-Galactose; α-D-Glucose; m-Inositol; D-Mannitol; D-Mannose; L-Rhamnose; D-Sorbitol; Methyl Pyruvate; Mono-Methyl-Succinate; Acetic Acid; Cis-Aconitic Acid; Citric Acid; Formic Acid; D-Galactonic Acid Lactone; D-Gluconic Acid; D-Glucosaminic Acid; D-Glucuronic Acid; α-Hydroxy Butyric Acid; β-Hydroxy Butyric Acid; p-Hydroxy Phenylacetic Acid; α-Keto Butyric Acid; α-Keto Glutaric Acid; D,L-Lactic Acid; Malonic Acid; Propionic Acid; Quinic Acid; D-Saccharic Acid; Sebacic Acid; Succinic Acid; Bromo Succinic Acid; Succinamic Acid; L-Alaninamide; D-Alanine; L-Alanine; L-Alanyl-glycine; L-Asparagine; L-Aspartic Acid; L-Glutamic Acid; Glycyl-L-Glutamic Acid; L-Histidine; Hydroxy-L-Proline; L-Leucine; L-Ornithine; L-Phenylalanine; L-Proline; L-Pyroglutamic Acid; D-Serine; L-Serine; L-Threonine; γ-Amino Butyric Acid; Urocanic Acid; Inosine; Uridine; Glycerol. The majority of characteristics for the type strain are in agreement with other six tested representatives of *B. agricolaris* sp. nov. *Burkholderia hayleyella* sp.nov. are susceptible to tetracycline at 30μg/ml.

#### Differences between *Burkholderia* sp.nov.

*B. agricolaris* BaQS159 is able to utilize N-Acetyl-D-galactosamine, N-Acetyl-D-glucosamine, Adonitol, L-Arabinose, D-Arabitol, D-Fructose, D-Galactose, m-Inositol, D-Mannitol, L-Rhamnose, D-Sorbitol, Mono-Methyl-Succinate, Cis-Aconitic Acid, Citric Acid, Formic Acid, D-Galactonic Acid Lactone, D-Glucosaminic Acid, D-Glucuronic Acid, p-Hydroxy Phenylacetic Acid, Malonic Acid, Quinic Acid, D-Saccharic Acid, Sebacic Acid, Succinamic Acid, L-Alaninamide, L-Alanine, L-Histidine, Hydroxy-L-Proline, L-Leucine, L-Ornithine, L-Phenylalanine, L-Pyroglutamic Acid, D-Serine, γ-Amino Butyric Acid, Urocanic Acid, Inosine, Uridine, and Glycerol which *B. hayleyella* BhQS11 cannot.

*B. agricolaris* BaQS159 is able to utilize N-Acetyl-D-galactosamine, Adonitol, L-Arabinose, D-Arabitol, L-Fucose, D-Galactose, m-Inositol, D-Mannitol, L-Rhamnose, D-Sorbitol, Mono-Methyl-Succinate, Acetic Acid, Formic Acid, D-Galactonic Acid Lactone, D-Glucuronic Acid, p-Hydroxy Phenylacetic Acid, Malonic Acid, Quinic Acid, D-Saccharic Acid, Sebacic Acid, L-Histidine, Hydroxy-L-Proline, L-Ornithine, L-Phenylalanine, L-Pyroglutamic Acid, D-Serine, γ- Amino Butyric Acid, Urocanic Acid, Uridine, and Glycerol which *B. bonniea* BbQS859 cannot.

*Burkholderia hayleyella* (hey′lee.el.uh. N.L. fem. adj. *hayleyella*, pertaining to Hayley. This specific epithet is in honor of the daughter of Debra A. Brock). Colony morphology is off-white, domed, and shiny with smooth edges. Bacteria are motile, non-sporulating, straight rods. The strains are stored in a sterile 20% glycerol solution at −80°C and subcultured on SM/5 agar plates at 22°C. We isolated the type strain BhQS11 as a symbiont of wild *D. discoideum* clone QS11 in February 2008. *D. discoideum* QS11 was isolated from soil and leaf litter collected from Mountain Lake Biological Station in October 2000. The G+C content is 59·24 mol% calculated from whole genomic sequence. Good growth at 30°C, weak growth at 22°C, and no growth at either 4°C, 37°C, or 45°C on nutrient agar containing glucose, bactopeptone, and yeast extract. The type strain BaQS11 has the ability to utilize the following carbon sources as determined by the Biolog GN2 test panel: Tween 40; Tween 80; L-Fucose; α-D-Glucose; D-Mannose; Methyl Pyruvate; Acetic Acid; D-Gluconic Acid; α-Hydroxy Butyric Acid; β-Hydroxy Butyric Acid; α-Keto Butyric Acid; α-Keto Glutaric Acid; D,L-Lactic Acid; Propionic Acid; Succinic Acid; Bromo Succinic Acid; D-Alanine; L-Alanyl-glycine; L-Asparagine; L-Aspartic Acid; L-Glutamic Acid; Glycyl-L-Glutamic Acid; L-Proline; L-Serine; L-Threonine. The majority of characteristics for the type strain are in agreement with other six tested representatives of *B. hayleyella* sp. nov. *Burkholderia hayleyella* sp.nov. are susceptible to 0.1μg/ml ampicillin plus 0.3 μg/ml streptomycin sulphate, and to tetracycline at 30μg/ml.

Differences between *Burkholderia* sp.nov.: *B. hayleyella* BhQS11 is able to utilize L-Fucose and Acetic Acid which *B. bonniea* BbQS859 cannot. *B. hayleyella* BhQS11 utilizes a smaller subset of the same carbons as *B. agricolaris* BaQS159.

*Burkholderia bonniea* (baan´-ee-uh. N.L. fem. adj. *bonniea*, pertaining to Bonnie. This specific epithet is in honor of the mother of Susanne DiSalvo). Colonies are off-white, shiny, and domed with smooth edges. Bacteria are motile, non-sporulating, straight rods. The strains are stored in a sterile 20% glycerol solution at −80°C and subcultured on SM/5 agar plates at 22°C. We isolated the type strain BbQS859 as a symbiont of wild *D. discoideum* clone QS859 in August 2014. *D. discoideum* QS859 was isolated from deer feces collected from Mountain Lake Biological Station in July 2014. The G+C content of the type strain is 58·7 mol% calculated from whole genomic sequence. Good growth at 30°C, weak growth at 22°C, and no growth at either 4°C, 37°C, or 45°C on nutrient agar containing glucose, bactopeptone, and yeast extract. The type strain BaQS859 has the ability to utilize the following carbon sources as determined by the Biolog GN2 test panel: Tween 40; Tween 80; N-Acetyl-D-glucosamine; D-Fructose; α-D-Glucose; D-Mannose; Methyl Pyruvate; Mono-Acetic Acid; Cis-Aconitic Acid; Citric Acid; D-Gluconic Acid; D-Glucosaminic Acid; α-Hydroxy Butyric Acid; β-Hydroxy Butyric Acid; α- Keto Butyric Acid; α-Keto Glutaric Acid; D,L-Lactic Acid; Propionic Acid; Succinic Acid; Bromo Succinic Acid; Succinamic Acid; L-Alaninamide; D-Alanine; L-Alanine; L-Alanyl-glycine; L-Asparagine; L-Aspartic Acid; L-Glutamic Acid; Glycyl-L-Glutamic Acid; L-Leucine; L-Proline; L-Serine; L-Threonine; Inosine.

#### Differences in carbon utilization between *Burkholderia* sp.nov.

*B. bonniea* BbQS859 is able to utilize N-Acetyl-D-glucosamine, D-Fructose, Mono-Acetic Acid, Cis-Aconitic Acid, Citric Acid, Succinamic Acid, L-Alaninamide, L-Alanine, L-Leucine, and Inosine which *B. hayleyella* BhQS11 cannot. *B. bonniea* BbQS859 is able to utilize Mono-Acetic Acid which *B. agricolaris* BaQS159 cannot.

## Discussion

We use several kinds of evidence to delineate *B. agricolaris, B. hayleyella*, and *B. bonniea* as new species. We have tested the closest 16S relatives and found only these three sp. nov. have the ability to colonize *D. discoideum*, to be carried through multiple amoebae to fruiting body cycles, and to facilitate the carriage of food bacteria to seed new environments (4). We place these new species in a phylogeny which shows they each comprise fully supported independent clades. Phylogenetic evidence also show *B. hayleyella*, and *B. bonniea* strongly diverged from other *Burkholderia* species. We analyzed physical and phenotypic traits of carbon usage, bacterial length, and optimal growth temperature and found significant differences from non-symbiont *Burkholderia*. The three *Burkholderia* sp. nov. have also diverged from each other in pairwise carbon usage. Moreover, both *B. agricolaris* and *B bonniea* are able to utilize some carbons that non-symbiont *Burkholderia* cannot. These data support the identification and naming of three new *Burkholderia* species.

Our three *Burkholderia* sp. nov. have a facultative endosymbiotic lifestyle with their host *Dictyostelium discoideum*. A common feature of endosymbiosis is the streamlining and loss of non-essential genes (9). Several lines of evidence suggest cell size corresponds positively with genome size. Examples are found in red blood cells and genome size of vertebrates where the red blood cell increases with genome size (10). Using avian genomes known to be small and streamlined compared to other vertebrates, Organ et al. found a similar pattern of correspondence between fossilized osteocytes and predicted genome size in extant vertebrates (11). Beaulieu et al. also found a similar pattern in a broad array of 101 angiosperms showing cell size and genome size scale positively, something that has proven generally true in plants (12) (13). Here, we demonstrate that the lengths of the symbiotic *Burkholderia* sp. nov. bacteria are smaller than their free-living closest relatives. Additionally, the type strains of *B. hayleyella, B. bonniea*, and *B. agricolaris* have lost the ability to utilize many of the 95 carbons tested compared to the non-symbiont *Burkholderia* tested suggesting corresponding gene losses based on loss of function. These data taken together suggest genome streamlining of non-essential genes for the three Burkholderia sp. nov. consistent with an endosymbiotic lifestyle.

Lastly, the optimal growth temperature of the three *Burkholderia* sp. nov. at 30°C is lower than the non-symbiont outgroup species at 37°C. One possible explanation could be that this is an adaptation to the much lower optimal growth temperature range of their host *D. discoideum* which is 20-25°C (14)

In sum, these three new species have diverged from their ancestors in measurable ways that are likely to be due to their endosymbiotic habit within *D. discoideum*. We suggest classifying these isolates as novel species for which the names *Burkholderia agricolaris, Burkholderia hayleyella*, and *Burkholderia bonniea* are proposed with the type strains BaQS159 (Dictybase DBS0351125; NCTC xx), BhQS11 (Dictybase DBS0351126; NCTC xx), and BbQS859 (Dictybase DBS0351127; NCTC xx) respectively, as the type strains.

## Materials and Methods

### Bacteria isolates

Wild *Burkholderia* symbiont strains used in this study were isolated from the Queller and Strassmann (QS) *D. discoideum* collection. See Supplementary Table 1 for collection locations and GPS coordinates. We previously isolated *Burkholderia* strains BhQS11, BhQS21, BhQS22, BhQS23, BhQS155, BaQS159, and BaQS161 (15) and *Burkholderia* strains BaQS70, and BaQS175 (4). We then isolated *Burkholderia* strains BaQS31, BhQS46, BhQS69, BaQS80, BhQS115, BaQS317, BbQS433, BhQS530, BbQS859, BaQS983, BaQS1007, and BaQS1045. We used *Burkholderia fungorum* ATCC BAA-463, *Burkholderia xenovorans* LB400, and *Burkholderia phymatum* STM-815 for our non-symbiont, close relative reference strains. See Supplementary Table 1 for specific strains used in preparing the phylogeny and for examining carbon usage, cell length, and optimal growth temperatures. The strains are stored in a sterile 20% glycerol solution at −80°C and subcultured on SM/5 agar plates (2 g glucose, 2 g Oxoid bactopeptone, 2 g Oxoid yeast extract, 0.2 g MgSO_4_, 1.9 g KH_2_PO_4_, 1 g K_2_HPO_4_, and 15.5 g agar per liter DDH_2_O) at 22°C. To propagate bacteria from frozen stocks for experimental assays, we plated on SM/5 nutrient agar plates and grew at 22°C to stationary phase.

### Phylogeny

We used k-mers, DNA “words” of a given length k, to estimate the phylogenetic relationships of the symbionts relative to other known plant-associated and environmental *Burkholderia*. Such alignment-or even genome assembly-free methods are increasingly available for many types of analyses that leverage next-generation sequencing data, including phylogenetic reconstruction (16). These types of methods are particularly powerful because they can combine assembled and unassembled genome sequencing data since k-mer frequencies can be estimated from either data source. These methods also improve upon the shortfalls of species delineation using 16S (17) as they can take into account sequence characteristics of entire genomes.

We first estimated phylogenies using three different k-mer sizes, k=23, 29, and 31 with AAF (Assembly and Alignment-Free) version 20160831 (18). The number of potential k-mers increases by a factor of 4 as k-mer size increases. The appropriate size of k-mer to adopt is thus a balance between information content (larger k-mers contain more information) and computational efficiency (smaller k-mers require less computation and memory). Because all topologies were identical for the k-mer sizes we tested, we chose k=23 for our final k-mer size. Next, we ran 999 nonparametric bootstraps and resampled 1/k rows (k=23) from the shared k-mer table as described by the developer of this software (https://github.com/fanhuan/AAF accessed Dec 2017). These bootstraps indicated that our topology was robust (all bootstrap values = 1).

### Carbon Usage

We used Biolog GN2 Microplates to determine carbon source usage patterns for each bacterial isolate (Biolog Inc., Hayward, CA). These plates contain 95 test carbons and one blank control well. The 95 test carbons correspond to 11 carbon groups such as carbohydrates and amino acids (see Supplementary Table 2 for complete list of carbon groups and individual carbons in each group). We brought the plates to room temperature prior to filling. We made suspensions in non-nutrient buffer for each bacterial isolate at OD_600_ 0.7. We then added 150uL of this bacteria suspension to each well of the GN2 plate. Plates were incubated at 30°C for 24 hours, at which point they were photographed and the optical density was measured using a Tecan Infinite M200Pro plate reader (Wavelength = 590nm, bandwidth = 9nm, 5 flashes per well). We scored the results from the Biolog tests binomially - either the well was positive, meaning that the bacteria could use the substance as a carbon source, or the well was negative and it was not an available carbon source. To determine if a well was positive or negative, we calculated if the absorbance of each well minus the blank control absorbance is above or below 97.5% of the blank control. This is equivalent to a 5% confidence for a two-tailed distribution. We determined the negative baseline independently for each plate based on the value of the blank control. We analyzed all data using R v3.4.1 (19) employing the following specific packages (20–22). We performed a principal component analysis on the ability of individual members of each *Burkholderia* sp. nov. and the three close sister *Burkholderia* to utilize carbon grouped into 11 carbon types (See Supplementary Table 2 for individual carbons in each group). To test the effect of carbon usage by *Burkholderia* species, we used a generalized linear mixed model with a random-slope and a binomial error distribution. We used *Burkholderia* clone as our random factor, with clade and carbon type as fixed factors, and ability to use a particular carbon source as the response. We also compared each carbon source between species pairs (Supplementary Table 3) with *post hoc* comparisons and Benjamini-Hochberg adjusted P-values (23).

### Bacterial Length

To examine morphological characteristics, we grew each bacterial isolate from frozen stocks on SM/5 agar plates (24) for about 4 days to stationary phase. We then collected and prepared a bacterial suspension of each test isolate in non-nutrient buffer (2.25 g KH_2_PO_4_ and 0.67 g K_2_HPO_4_ per liter DDH_2_O) at OD_600_1.5. To prepare fixed bacteria for imaging by microscopy, we first prepared the fixative solution by adding 6.26 μL of 8% gluteraldehyde per one mL of 16% paraformaldehyde (Electron Microscopy Sciences, Hatfield PA USA). Next, we added 200μL of each bacterial suspension, 8μL of 1M NaPO_4_ pH 7.4, and 40μL of the fixative solution to a 1.5ml. Eppendorf tube and gently mixed. We incubated the reactions for 15 minutes at room temperature, followed immediately by 30 minutes on ice. We then centrifuged briefly at 10,000 g to pellet and wash the bacteria, repeating 3 times using phosphate buffered saline (PBS; Fisher Scientific, Pittsburg PA, USA), and ultimately resuspending in 1ml PBS. We prepared the microscope slides for image capture by adding 200μL of 1% agarose in PBS (melted and slightly cooled) onto a single-depression microscope slide (VWR, Radnor PA, USA) and immediately overlaid with a cover slip. After 10 minutes of cooling, we removed the cover slip and added 5μL of the fixed bacteria samples directly onto the agarose pad. Once the bacteria solution dried, the coverslip was replaced. We captured images and measured the lengths of about 100 individual bacteria for each isolate using a Nikon TI-E microscope and NIS-Elements software (see Supplementary Table 4 for exact number measured for each isolate). Using R and package (21), we compared bacterial lengths among each isolate using a generalized linear mixed model with a random-slope and a negative binomial error distribution. We used *Burkholderia* clone as our random factor and clade as a fixed factor, with cell length as the response. We made *post hoc* comparisons to test the effect of differences in bacterial length between species pairs using Benjamini-Hochberg adjusted P-values (23).

### Optimal growth temperature

To determine the temperature for optimal growth for each species, we streaked all clones on SM/5 plates and placed them in 5 different temperatures (4°C, 22°C, 30°C, 37°C, and 45°C). We examined the plates and photographed them at 24 hours. We scored plate growth using the following categories: no growth, little growth, moderate growth, and excellent growth. We recorded optimal growth temperature for each isolate based on the temperature at which the bacteria grew the most densely. We made a heat map of the data (Figure 4) and performed a Fisher’s Exact Test with a 4 × 3 matrix testing the 22°C, 30°C, and 37°C temperature data to look for correlations (22). We excluded 4°C and 45°C data from the analysis because none of the isolates including the controls grew at these temperatures.

### Data deposition

All data are deposited in the Dryad Digital Repository (will deposit after acceptance).

## Author Contributions

DB, AH, SN, SD, TH, DQ, and JS conceived of the study. AH and DB performed the experimental assays. SN constructed the phylogeny. KG did and advised on much of the statistical analyses. DB, AH, SN, KG, DQ, and JS wrote the final draft. All authors reviewed and approved the final draft.

## Acknowledgments

We thank the Queller/Strassmann lab group for much useful advice. We particularly thank Jason Zuke for help with microscopy and Joe LaManna for help with principal component analysis in R. We are grateful to James Tiedje at Michigan State University for *Burkholderia xenovorans* LB400 and to Dr. Lionel Moulin at IPME, University of Montpellier, France for *Burkholderia phymatum* STM 815.

